# Genotyping Test Development and Genotyping Survey of Pakistani population of Holstein Friesian imported from Different Origins for A1/A2 SNP in Beta-casein Gene

**DOI:** 10.1101/720045

**Authors:** Waqas Rafique Ali, Imran Amin, Muhammad Asif, Shahid Mansoor

**Author notes:** Corresponding Author: Shahid Mansoor.

## Abstract

Quality of milk is determined by multiple factors including microbial load, somatic cell score, content percentages and absence of adulterants. However, in term of genetics, milk quality can be linked with beta casein. Beta casein is a milk protein having genetic variants, one of which is believed to be linked with progression of many diseases including cardiovascular diseases, metabolic diseases and nervous disorders. As beta casein has genetic basis so milk quality can be elucidated by animal genotyping. But no such genotyping facility was available country wide for this purpose. Hence objective of this study is the development of diagnostic test using ARMS-PCR technology in Pakistan and genotyping of beta casein variants in Pakistani population of Holstein Friesian cattle. A total of 166 blood samples were collected from Holstein Friesian (imported from Holland, USA and Australia) and genotyped using ARMS-PCR. Validation of ARMS-PCR was confirmed by Sanger sequencing of beta casein gene from 78 animals. ARMS-PCR based test was then compared with parallel genotyping techniques in term of cost and time effectivity. Reproducibility of this test was tested and validation of this test was done by Sanger sequencing. This test is cost effective as well as time effective in compared to other techniques. Holstein Friesian originating from different countries have different percentages of beta casein variants. This test can be used routinely for genotyping of beta casein. In future there is a need to find out prevalence of beta casein variants of indigenous cattle as well as cross bred population of Pakistan and formulation of breeding policy, keeping in view health concerns of A1 genotype of β-casein.

**Novelty Statement:** In this study, a diagnostic test is developed to genotype beta casein variants in animal. This is the first report of its kind in Pakistan as before this report either sequencing or tedious and costly procedures were in use for genotyping of beta casein variants in Pakistan. Additionally, for the first time in Pakistan Holstein Friesian imported from Holland, United States and Australia were genotyped to elucidate their genotype status for beta casein polymorphism.

## Introduction

Milk is rich source of nutrients and has remained important part of diet in history. In ancient era of Egypt milk and dairy products were considered as specialty and were reserved for upper class only including royal families and wealthy people. By 5^th^ century, in Europe, cows and sheep were prized on the basis of milk production. European dairy was then brought to north America in early 1600’s where revolution in milk consumption happened when pasteurization was introduced which enabled the use of milk by distribution away from farm keeping quality undeteriorated to stipulated time. Now this has become need of every home with estimated per capita consumption of 20 liters in north America. In spite of established quality standards for microbial load, somatic cells count, proximate composition, there are other factors too which have not been under limelight until publishing of book titled “Devil in the Milk: Illness, Health and the Politics of A1 and A2 Milk” (Woodford 2009). In this book details of beta casein variants and their effects on human health has been hypothesized. This started an uprising global debate over milk quality with respect to beta casein and its relevance with disease occurrence.

Milk contains 3.2 % of protein by composition and in proteins about 80% is casein which exists in milk in the form of micelles. Remaining 20% of total protein is called whey protein which comprises of α-lactalbumin and β-lactoglobulins. Casein class of proteins further have many types, α-casein, β-casein and κ-casein (Jones 2002). β-casein protein constitutes 25-30% of total milk protein. Beta casein protein is encoded by *CSN2* gene that spans over 8.5 kb of chromosome number 6, comprising of 9 exons (Bonsing et al. 1988). A SNP believed to be associated with human health exists in exon 7 of *CSN*2. This SNP is generally named as A1A2 SNP. Difference between A1 and A2 variant is the substitution of cytosine (C) nucleotide with adenine (A) nucleotide at specific position, which leads to a change in the amino acid from histidine (present in A1 type) with proline (present in A2) at 67^th^ position of beta casein polypeptide (Bonsing et al. 1988).

Based upon the reports it is believed that consumption of A1 milk is linked with occurrence of many diseases which includes type 1 diabetes, cardiovascular diseases, autism, schizophrenia and auto immune disorders (Woodford 2008). A large-scale study, conducted using data from 19 countries in which a correlation between A1 casein and disease prevalence was analyzed concluded with the fact that countries having high per capita consumption of A1 milk have higher increased disease prevalence especially ischemic heart disease (Laugesen and Elliott 2003). This may be because A1 milk casein is subjected to 3-dimensional conformational change in protein structure which makes it vulnerable to different proteases especially digestive enzymes (Figure 3). In *in vitro* studies, pepsin, elastase, leucine aminopeptidase (LAP) and pancreatin are found to be able to digest beta casein protein. However, this enzymatic action leads to the release of an oligopeptide of 7 amino acids named β-caseomorphine 7 (BCM-7) while this opioid agent is not released by A2 β-casein variant (Jinsmaa and Yoshikawa 1999).

**Figure 1:**
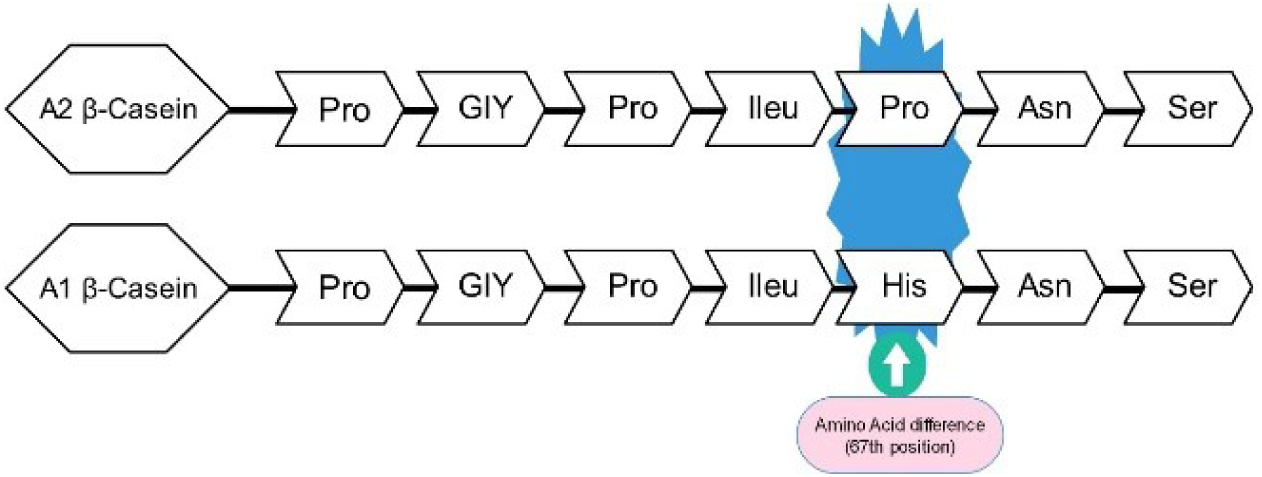
Replacement of Proline amino acid (A2 beta casein) with histidine amino acid (A1 beta casein) at 67th position of beta casein protein due to SNP

**Figure 2:**
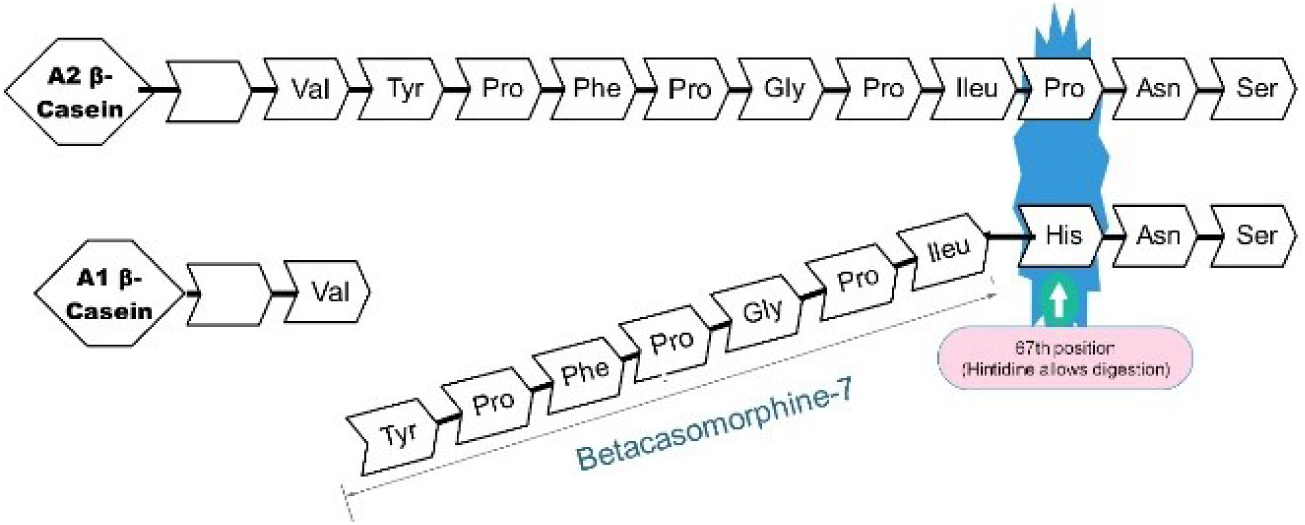
Production of betacaseomorphine-7 (BCM-7) as a result of catalytic activity of proteolytic enzymatic action due to conformational change of A1 beta casein. A2 beta casein variant do not yield BCM-7 due to conformational change.

**Figure 3:**
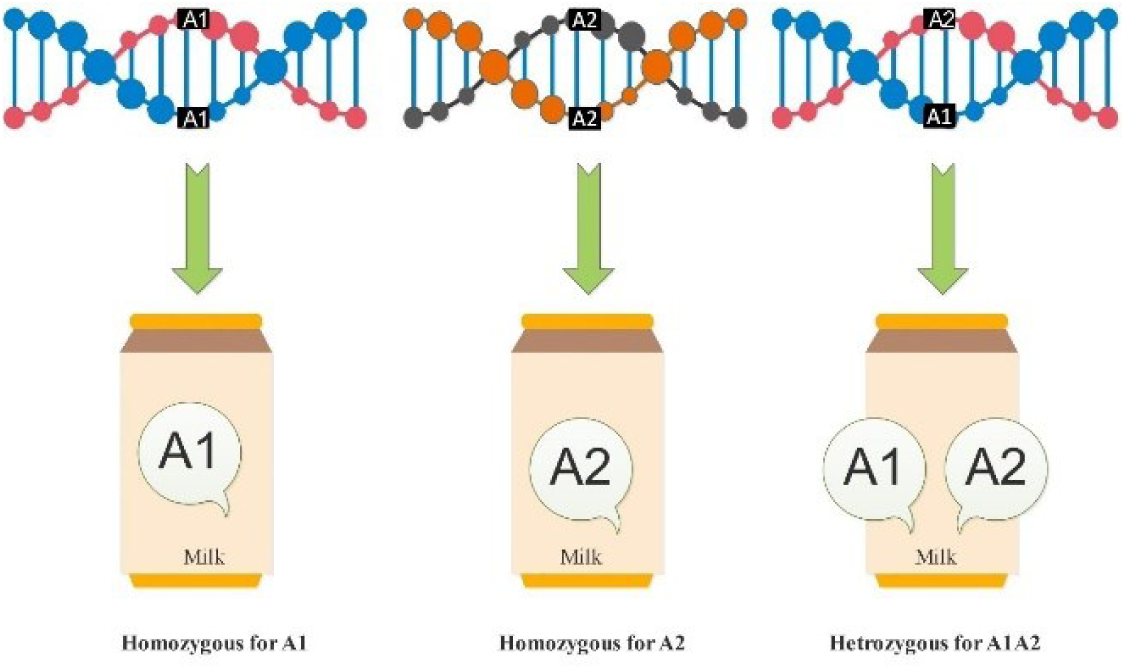
Co-dominant expression of CSN-2 gene. A1A1 genotype produces A1 beta casein in milk. A1A2 genotype produces both A1 and A2 beta casein protein in milk and A2A2 genotype produces A2 beta casein protein in milk

It is believed that A1 type of beta casein is secreted in milk by European cattle while cattle of Asian or African origin secretes A2 type of beta casein in milk, also animals other than cattle produce A2 milk. Even human β-casein gene sequence shows that it contains A2 milk (Ng-Kwai-Hang and Grosclaude 2003). It is worth mentioning that A1A2 is co-dominant in expression, which means that if animal is heterozygous it would produce both milk casein types in milk. There is no effect of dominancy of one allele over other (Fig.3).

Pakistan ranks at number six in terms of number of people with diabetes worldwide. Moreover, 30 to 40 percent of all deaths are due to cardiovascular diseases (CVD) in Pakistan (https://www.shifa.com.pk/chronic-disease-pakistan/). Keeping in view the public health importance and alarming facts of disease prevalence there is a need of technology to test the genotype of livestock of population of Pakistan. There are two approaches to test beta casein variant in milk. (1) Milk can be tested for the presence of A1 and A2 variant of beta casein in milk, but it requires so much effort to elucidate the conformational change between two proteins of same length, proteomics study is cumbersome and requires sophisticated instruments. (2) Animals are tested at genetic level, as this trait is co-dominant in expression hence milk produced from and A1 animal will contain A1 protein only, milk produced from A2 genotype animal will produce A2 milk only and animal of heterozygous genotype will produce both A1 and A2 variant of beta casein.

No indigenous technology was available in Pakistan to genotype animals for the elucidation of milk status in term of beta casein variant. In Pakistan, there are some studies on the beta casein variants in bovine but either these studies did not involve A1A2 SNP (Hamza, Wang, and Yang 2010; Mir, Tahir, and Sheikh 2013) or they directly sequenced the portion of gene instead of development of technology for genotyping (Shahla 2016). Some international companies provide A1A2 genotyping services. Companies includes CLARIFIDE™, GeneMark®, Veterinary Genetics Laboratory in UC Davis-USA, ICAR-NBAGR-India, GENESEEK-USA and others. Different technologies are employed for this test in different labs which mainly incudes allele-specific PCR, amplification created restriction site PCR, single stranded conformation polymorphism PCR (SSCP-PCR), Taq Man method, liquid chromatography– mass spectrometry (LC-MS) and DNA sequencing. Each technique has its own pros and cons.

SNP genotyping technique used in this study is allele specific PCR called Amplification Refractory Mutation System coupled with PCR (ARMS-PCR) (Medrano and de Oliveira 2014). In this technique four primers are used hence called tetra primer ARMS-PCR. It is found more convenient, time effective, cost effective and relatively simple method to genotype animals for A1A2.

According to a rough estimate, Pakistan is believed to have 10 million heads of Holstein Friesian and major part of this is contained in corporate dairy sector, milk of which is processed and marketed across the country. Hence there is a need of survey to find genotype status of imported animals. At present there are three major sources of import of these animals viz America, Australia and Holland. It was seen that no proper certification is provided at the time of import of cattle regarding genotype of beta-casein polymorphism. Hence it was important to know the genotype of these animals that may influence the policy making.

## Materials and Methods

A test population comprising of 78 Holstein cattle were selected for development ARMS-PCR based genotyping test. Sampling of test population (n=78) was done from a farm situated in Renala Khurd Tehsil of district Okara, Punjab. After optimization of genotyping test, blood samples of Holstein Friesian cattle were drawn that were imported from different countries of world for the purpose of genotype survey of β-casein gene polymorphism. Details of sampling places is shown in figure (Map). Twenty-five (25) blood samples were of Holstein Friesian imported from Dutch countries, 36 were of American origin and 27 were of Australian origin. All animals were female. Complete randomness is ensured in selection of farms and animals for blood sampling. All farms belong to private sector. All animals were in lactation. Blood samples were collected form jugular vein after proper restraining of animals and by following strict hygienic guidelines under direct supervision of licensed veterinary practitioner. Blood was collected in BD Vacutainer™ Hemogard Closure Plastic K3-EDTA Tube. After proper mixing of blood with K3-EDTA, vacutainers were kept in icebox for transportation. Upon arrival in laboratory, DNA was extracted using FavorPrep™ blood genomic DNA extraction mini kit. Integrity of DNA was checked by electrophoresis using 1% agarose gel. Quality and quantity of DNA was determined by Nanodrop® 2000C Spectrophotometer.

Primers were designed by a tool named primer1 (Collins and Ke 2012) and synthesized commercially by Eurofins Genomics® Pvt. Ltd. Sequences of primers are given in table 2.

**Table 1:**
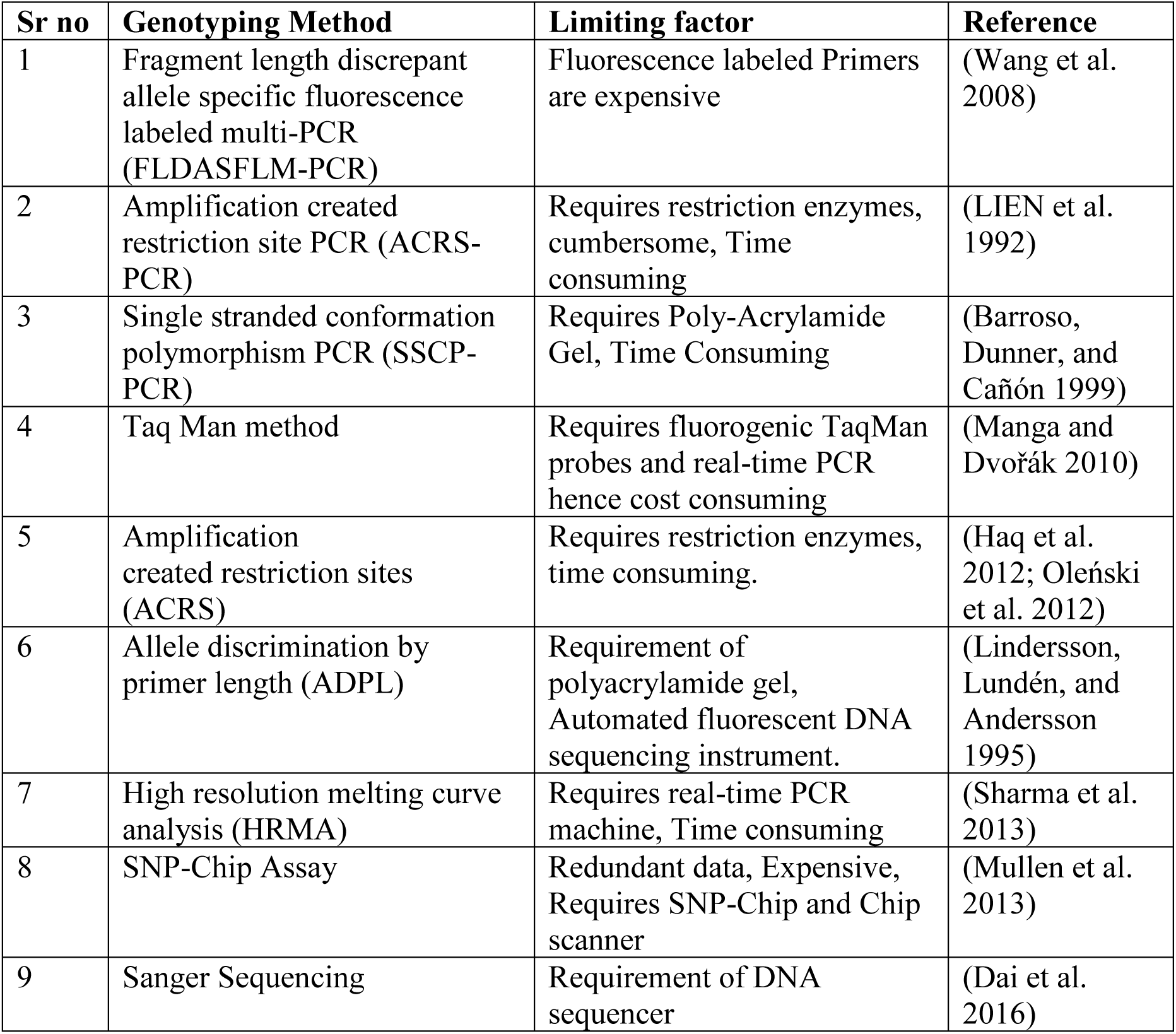
Comparison of genotyping methods with special reference to bovine beta casein A1A2 SNP

| Sr no | Genotyping Method | Limiting factor | Reference |
| --- | --- | --- | --- |
| 1 | Fragment length discrepant allele specific fluorescence labeled multi-PCR (FLDASFLM-PCR) | Fluorescence labeled Primers are expensive | (Wang et al. 2008) |
| 2 | Amplification created restriction site PCR (ACRS-PCR) | Requires restriction enzymes, cumbersome, Time consuming | (LIEN et al. 1992) |
| 3 | Single stranded conformation polymorphism PCR (SSCP-PCR) | Requires Poly-Acrylamide Gel, Time Consuming | (Barroso, Dunner, and Cañón 1999) |
| 4 | Taq Man method | Requires fluorogenic TaqMan probes and real-time PCR hence cost consuming | (Manga and Dvořák 2010) |
| 5 | Amplification created restriction sites (ACRS) | Requires restriction enzymes, time consuming. | (Haq et al. 2012; Oleński et al. 2012) |
| 6 | Allele discrimination by primer length (ADPL) | Requirement of polyacrylamide gel, Automated fluorescent DNA sequencing instrument. | (Lindersson, Lundén, and Andersson 1995) |
| 7 | High resolution melting curve analysis (HRMA) | Requires real-time PCR machine, Time consuming | (Sharma et al. 2013) |
| 8 | SNP-Chip Assay | Redundant data, Expensive, Requires SNP-Chip and Chip scanner | (Mullen et al. 2013) |
| 9 | Sanger Sequencing | Requirement of DNA sequencer | (Dai et al. 2016) |

**Table 2.**
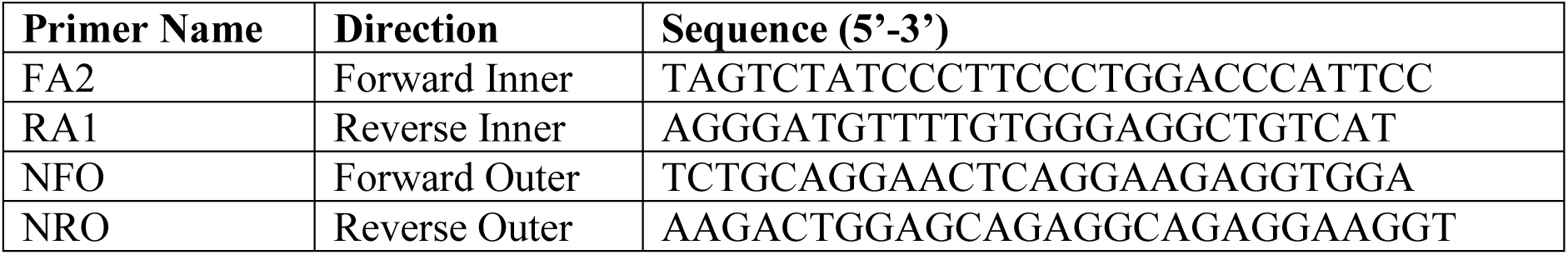
Primers for beta-casein genotyping.

Functionality of primers was checked pair wise, by *in silico* PCR tool hosted by GB-Shape website (Chiu et al. 2014). Primers were reconstituted and then diluted to the concentration of 100 pmol/μL. PCR was carried out using thermocycler c1000 Bio-Rad®. PCR was optimized using different annealing temperatures, primer concentrations, DNA concentrations and outer versus inner primer ratio. DreamTaq™ Green PCR Master Mix by Thermo Scientific™ was used in PCR. After development of test and checking for reproductivity, test was further validated by Sanger sequencing. Then using developed test, genotyping of Holstein Friesian of different origins was carried out.

## Results

PCR was optimized in 20 µL PCR reaction volume using recipe: Distilled water 5 µL, PCR Mastermix 10 µL, FA2 primer 0.75 µL, RA1 primer 0.75 µL, NFO primer 0.75 µL, NRO primer 0.75 µL, DNA Template 2 µL.

PCR was optimized at annealing temperature of 66 °C, DNA concentration between 40-100 ng/µL and outer versus inner primer concentration ratio of 1:1.

Optimized PCR profile is: Initial denaturation at 95°C for 5 mins, Cyclic denaturation at 94 °C for 1 min, primer annealing temperature 66 °C for 30 seconds, cyclic extension at 72 °C for 1 min and final extension at 72 °C for 10 mins with 34 amplification cycles.

PCR was carried out on test population comprising of 78 animals of Holstein Friesian breed. Most of these animals were of Australian origin. Amplified PCR products were visualized by electrophoresis using 1.5% agarose gel prepared in TAE buffer stained with ethidium bromide. Also, no-template control (NTC) was run in PCR to check contamination. A control band of 586 bp appeared in all samples while 384 bp fragment amplified in sample having A1A1 genotype, 256 bp amplified in sample having A2A2 genotype and both bands of 384bp and 256 bp appeared in animals that were in heterozygous state i.e. A1A2 as shown in figure 4.

**Figure 4.**
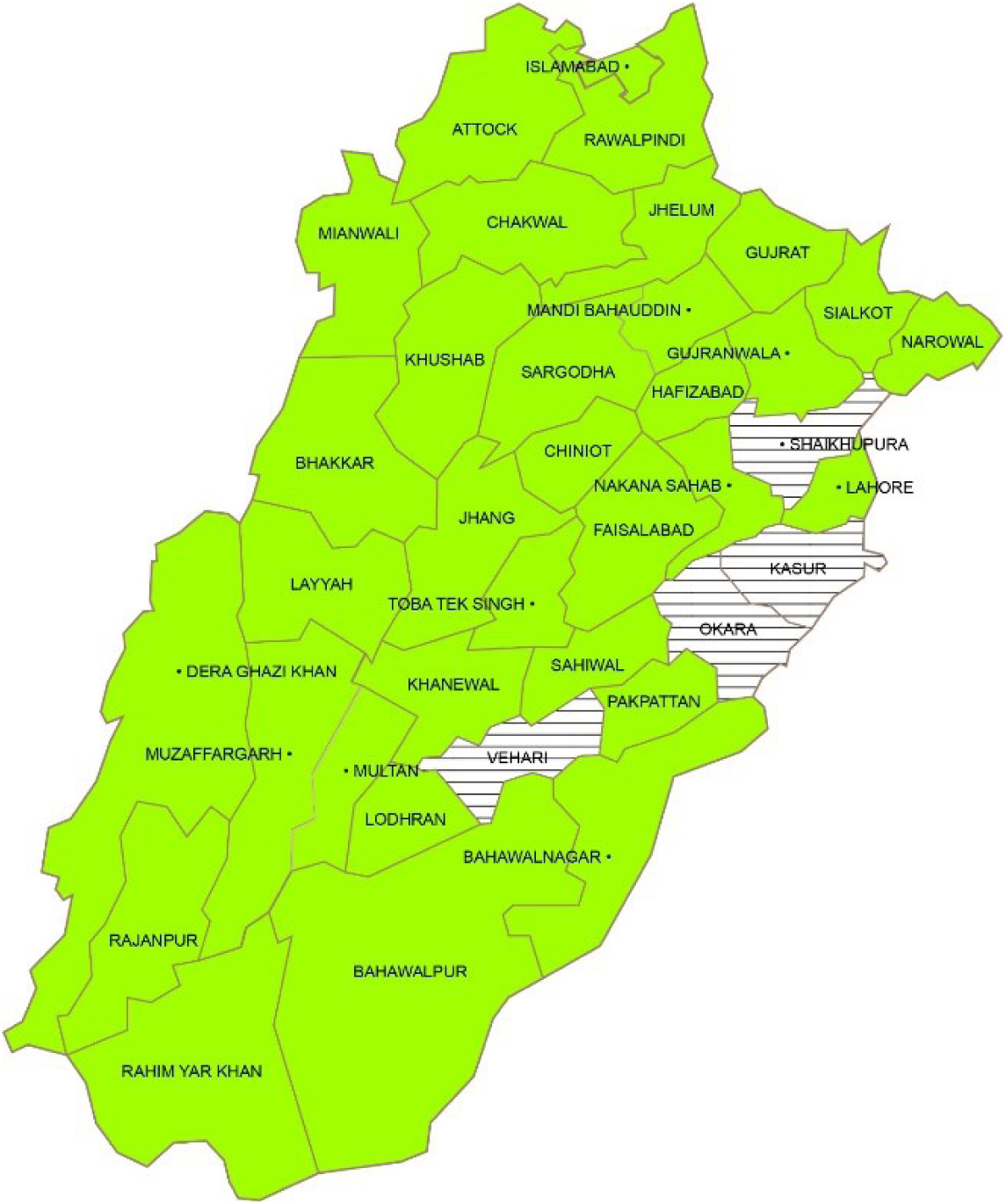
Map of province Punjab, Pakistan, Shaded area shows sites of sample collection for bovine beta casein genotyping studies.

Out of 78 samples of test population, sequencing results showed that 9 samples were having “A” allele at SNP position, 34 DNA samples were having both “A” and “C” allele and 35 samples were containing “C” allele. Samples having only “A” allele are homozygous for A1, containing both “A” and “C” allele are heterozygous i.e. A1A2 and containing only “C” allele are homozygous for A2. Computed percentages are 12 % for A1A1, 45 % for A1A2 and 44 % for A2A2. Results of sequencing are in complete accordance with that of ARMS-PCR as shown in figure 5.

**Figure 5:**
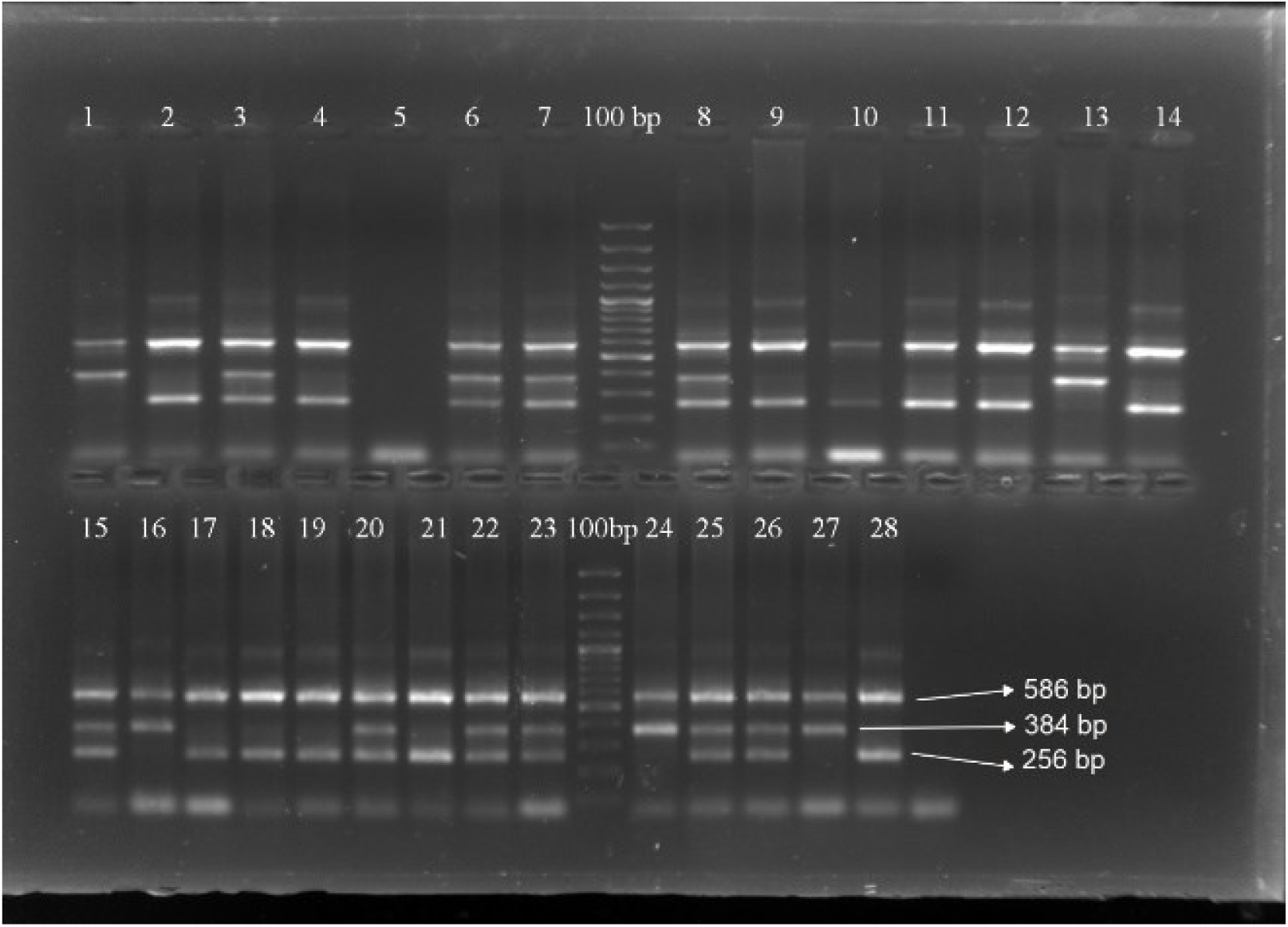
Gel picture of ARMS-PCR of bovine beta casein gene with 100 bp DNA marker. Band size of 384 bp shows A1 genotype and band size of 256 bp shows A2 genotype. Genotype of sample in lane having only 384 bp band is A1A2, that of only 256 bp is A2A2 and containing both 384 bp and 256 bp is heterozygous i.e. A1A2.

**Figure 6.**
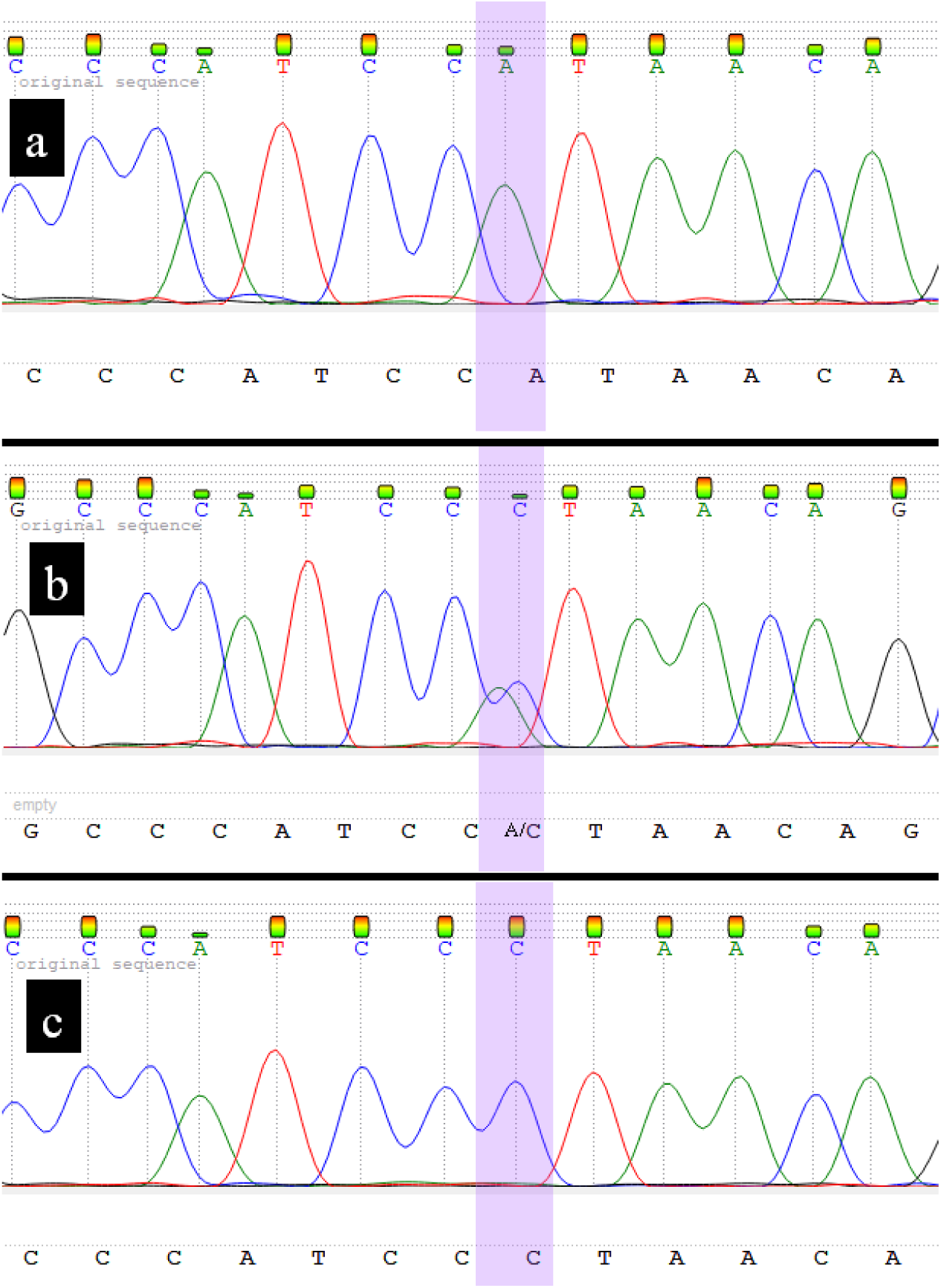
Sequencing Picture of fragment of beta casein gene. In sequence chromatogram A, there is A nucleotide in SNP position hence animal genotype is A1A1, In sequence chromatogram B, there are two peaks at SNP position, one for A and one for C, hence this animal is hetrozygus (A1A2), In chromatogram C, there is peak of C nucleotide at SNP position hence animal genotype is A2A2.

Genotyping survey of Holstein Friesian of different origin demonstrated that Holstein Friesian of Australian origin are found to be 18.4% A1A1, 52.2% A1A2 and 29.4% A2A2, Holstein Friesian imported from the States were 17% A1A1, 33% A1A2 and 50% A2A2 while Holstein Friesian of Dutch origin were found to be 12% A1A1, 40% A1A2 and 48% A2A2.

## Discussion

AMRS-PCR have been used for the SNP genotyping of different genes in different species including IL-10, TNF-α, TNF-β,TGF-β1 (Perrey et al. 1999), JAK2^V617F^ mutation in leukemia (Vannucchi et al. 2006), SNP genotyping in barley (Chiapparino, Lee, and Donini 2004) and other scores of SNPs in different species. There are other methods of SNP genotyping but AMRS-PCR has gained success due to its simplicity, cost effectivity, reproducibility, time effectivity and versatility. For high throughput genotyping advanced technologies like SNP chips can be utilized that has capability of genome wide genotyping but such technologies require complete setup. As TaqMan genotyping system has also got fame in recent years due to its high throughput nature, time effectivity, precision and validity, but it also requires labelled probes and sophisticated detection system (i.e. ABI Prism 7900HT system) for genotyping (Borodina, Lehrach, and Soldatov 2004; Giles et al. 2004; Hampe et al. 2001). Side by side comparison of different techniques for SNP genotyping has been discussed in table 1.

Genotyping of A1A2 beta casein in bovine has not been studied before in Pakistan using ARMS-PCR. There are few similar studies related to bovine genotyping. A study published in 2013 used tri-ARMS PCR for genotyping and they studied some variants of beta casein but not A1 and A2 variants (Mir, Tahir, and Sheikh 2013). In an Indian study, PCR-RFLP methodology is used to genotype variants of kappa casein and beta-lactoglobulin (Patel et al. 2008).

Genotyping survey showed that there is quite high percentage of Pakistani population of Holstein Friesian imported from different countries containing A1 allele. Due to which there are high chances of replacement herd having A1 genotype if this factor is not kept in consideration. However, by choosing selective breeding program, A1 allele can be diminished over time. It is obvious from results that Holstein cattle of Australian origin exhibit highest percentage of A1 homozygous animals and cattle of American origin exhibit highest percentage of A2 homozygous cattle. Similar studies have been conducted in past to find frequency of different genotypes of β-casein present in different cattle breeds in various countries. A study on Polish Holstein Friesian revealed 12.8% A1 homozygous cattle, 44.1% in heterozygous cattle (A1/A2) and 48.1% in A2 homozygous state (Olenski et al. 2010). Results of study conducted in this paper is in accordance with previous studies with minor variations. Keeping in view the prevalence of A1 genotype in Holstein Friesian present in Pakistan and health implications of milk from animals of A1 genotype, selective breeding program is suggested. Moreover, it is suggested that cattle and semen imported from different countries should be checked before use or certificate of genotype of animal or semen should be demanded from exporting country. As this test perform equally best for genotyping of indigenous cattle so in future there is need to genotype our local cattle population for the formulation of public policy regarding this issue.

